# Comparative Analysis of Antioxidant Activity, ROS, and Relative Water Content Between Red and Green Cabbage

**DOI:** 10.1101/2024.07.27.605406

**Authors:** Md. Rayhan Sojib, Nusrat Jahan Methela, Biswajit Das, Md Nure Adil Siddique, Sadia Sultana, Ataul Karim, Rafiqul Islam, Md Jahid Hasan Jone

## Abstract

Cabbage, the second largest leafy vegetable, is highly valued for its nutritional richness and versatility. As health consciousness increases, the worldwide demand for cabbage continues to grow steadily. Cabbages come in various forms, varying in size, color, texture, and nutritional properties. An experiment was conducted to distinguish significant differences in relative water content (RWC of leaf and RWC of head), relative oxygen species (MDA and H_2_O_2_), and antioxidant properties (POD, APX, and CAT) between red and green cabbage varieties. Cabbage samples were grown under fertilizer and control conditions to observe the impact of fertilizers on the acquisition of these properties. The results indicated that fertilizer application positively influenced the acquisition of relative water content, relative oxygen species, and antioxidant properties in both cabbage varieties. The results emphasized that red cabbage excelled in antioxidants and ROS levels, containing higher amounts compared to green cabbage. Conversely, green cabbage showed greater relative water content in both cultivation conditions. These findings suggest that consumers seeking higher antioxidant and ROS levels in their diet may benefit from incorporating more red cabbage into their meals. Further research into the mechanisms behind differences in red and green cabbage could inform breeding programs, enhancing nutritional traits for agricultural and dietary purposes.

## Introduction

Cabbage (*Brassica oleracea* var. *capitata*, 2n = 18), one of the most important cruciferous vegetables, is commonly grown worldwide (Singh et al. 2006) for its significance due to its high nutritional properties and wider adaptability. Due to improved livelihood and health consciousness, the requirement for vegetables and vegetable products is increasing day by day (Braga, Coletro, and Freitas 2019). Cabbage can be consumed either raw or processed in different ways, e.g., boiled, fermented, or used in salads, so a steady supply is required year-round. In 2020, world production of cabbages (combined with other brassicas) was 71 million tons (FAOSTAT 2022).

A wide range of vitamins, minerals, fiber, and carbohydrates can be obtained from the Cole crops which also contain high amounts of vitamin C, soluble fiber, and multiple potential anti-cancer nutrients (Chun et al. 2004). Among them, the antioxidant property is the most important dietary point of view. Phytochemicals such as vitamin C, flavonoids, carotenoids, and other phenolics are the constituents that contribute to this property. Due to its antioxidant (Vega-Galvez et al. 2023; Liang et al. 2019) anti-inflammatory (Kasarello et al. 2022), and antibacterial (Arrais et al. 2022) properties, cabbage has widespread use in traditional medicine (Gruszecki et al. 2022), in the alleviation of symptoms associated with gastrointestinal disorders (gastritis, peptic and duodenal ulcers, irritable bowel syndrome) as well as in the treatment of minor cuts and wounds and mastitis (Melim et al. 2022; Abramowitz et al. 2022; Tsania, Willy, and Sawitri 2023). Fresh cabbage juice, prepared either separately or mixed with other vegetables such as carrots and celery, is often included in many commercial weight-loss diets (Šamec et al. 2011).

The main varieties of cabbage cultivars are red, white (including spring greens and green), and savoy cabbages (Pérez and Cartea 2008). The different cultivated types of cabbage show great variation in terms of size, shape, color of leaves, and the texture of the head (Moreb et al. 2020; Singh et al. 2006). The mineral, nutritional, and antioxidant content varies throughout cabbage cultivars (Yue et al. 2024). Consequently, to maintain a balanced diet, understanding the distinctions between the major varieties of cabbage is crucial (Turner, Luo, and Buchanan 2020). Green cabbage (*Brassica oleracea* var. *capitata*) is the most commonly consumed type of cabbage. Previous research indicates that Red cabbage (*B. oleracea* var. *capitata* f. *rubra*) exhibits a superior nutritional profile, characterized by elevated levels of nutrients and bioactive compounds, thereby establishing its precedence over Green cabbage. Red cabbage boasts a wealth of nutrients, including amino acids and organic acids, making it a notable source of dietary fibre for individuals (Zayed et al. 2023). Recent investigations have further illuminated its polyphenolic content, particularly anthocyanins, renowned for their potent antioxidant properties and adeptness in scavenging free radicals (Ghareaghajlou, Hallaj-Nezhadi, and Ghasempour 2021). Epidemiological inquiries have also implicated red cabbage consumption in mitigating the risk of various ailments, encompassing cancer, cardiovascular disease, metabolic disorders, and Alzheimer’s disease (Manchali, Chidambara Murthy, and Patil 2012). Collectively, these findings underscore the favorability of red cabbage in dietary contexts, owing to its low-calorie profile, ample fiber content, and nutritional richness.

Prior studies (Ashfaq 2018; Yue et al. 2024) compared red and green cabbage according to quality indices and mineral elements; however, the antioxidants have not yet been examined. The present study aims to examine the antioxidant, reactive oxygen species, and relative water content of red and green cabbage as well as to determine how fertilizer affects these variables. This research might help in designing future breeding programs to meet nutritional needs through high-quality cabbage production.

## Materials and Methods

### Experimental site

The field experiment was conducted at the research field of the Noakhali Science and Technology University, Noakhali-3814, Bangladesh (22.79007, 91.10147). The annual average temperature of the region is 25.6 □ with an average rainfall of 3,302 mm, soil pH of 7.5, and salinity of 4.32 dSm^-1^ (Misu et al. 2023; Osman et al. 2020; Ahmed et al. 2024; Islam et al. 2023).

### Plant Materials and Experimental Design

Seedlings of two cabbage genotypes, Ruby King (Red cabbage from Takii and Co. Ltd. Japan) and Dream Ball (Green cabbage from East Bengal Seed Company, Bangladesh), were collected from Nobogram Khamarbari, Noakhali Sadar for the experiment. A Randomized Complete Block Design (RCBD) with three replications was used to conduct the experiment. Two treatments (fertilizers and genotypes) were used for this research and the applied fertilizer dosage rates were presented in the Supplementary Table.

### Sample preparation

A small piece of 0.1 g sample was taken from the head of each germplasm and stored in an icebox to prevent microbial growth and slow down enzymatic activity. These prepared samples were then examined to determine the relative water content, and reactive oxygen species (ROS) content and to measure their antioxidant activity. The biochemical analysis of the samples was conducted in the Department of Genetics and Plant Breeding, Bangladesh Agricultural University, Mymensingh-2202.

### Determination of Relative Water Content (RWC)

Fresh samples of leaf and head were collected, weighed, and dried in an oven at 70°C for 48 hours. The fresh and dry weight of the samples were used to determine the Relative Water Content using the following formula (González and González-Vilar 2003).

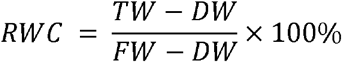

Here, TW = Total Weight; DW = Dry Weight and FW = Fresh Weight

### Determination of Reactive Oxygen Species Malondialdehyde (MDA) Activity

A 0.1g sample was ground with 2 ml of 0.1% TCA solution and centrifuged at 11000 rpm for 20 min. The supernatant was transferred to a new tube, mixed with 4ml of 20% TCA containing 0.5% TBA, and boiled at 70°C for 30 min before quickly cooling on ice. The resulting supernatant was collected after centrifugation at 5000 rpm for 5 min. Absorbance was measured at 532nm using a spectrophotometer, with all steps conducted at 4°C except for absorbance measurement. The following equation is used to calculate MDA concentration (Bispo et al. 2014).

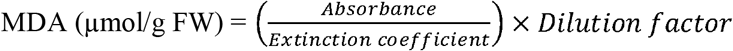

Here, Extinction coefficient of MDA = 155m/M/cm

### Hydrogen Peroxide (H_2_O_2_) Activity

A 0.1g sample was mixed with 2 ml of 0.1% TCA in a mortar and pestle at 4°C. After centrifugation at 11000 rpm for 20 min, the supernatant was transferred to a new tube and combined with phosphate buffer (10 mM) and potassium iodide (1M) in a 0.5 ml: 0.5 ml: 1 ml ratio, left in darkness for 1 hour. The absorbance was measured at 390 nm.

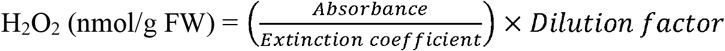

Here, Extinction coefficient of H_2_O_2_ = 0.28 µ/M/cm

### Determination of antioxidant activity

0.05 grams of each sample were homogenized with 3 mL of 50 mM potassium phosphate buffer (pH 8.0) and centrifuged at 11000 rpm for 20 minutes. The clear supernatant was then used to assay POD, APX, and CAT activity.

### Peroxidase activity (POD)

Peroxidase activity was assessed following the standard protocol (Nakano and Asada 1981). A mixture comprising 1.8 mL of 50 mM potassium phosphate buffer (pH 8.0), 0.3 mL of EDTA solution (2.5 mM), 0.3 mL of hydrogen peroxide (H_2_O_2_) solution (200 mM), and 0.3 mL of guaiacol solution (100 mM) was prepared in an Eppendorf tube. The enzymatic reaction commenced upon the addition of 0.1 mL of enzyme extract, and changes in absorbance were promptly recorded at 470 nm at 30-second intervals over a span of 2 minutes. Peroxidase activity was quantified based on the rate of absorbance increase per minute.

### Ascorbate Peroxidase Activity (APX)

Following the protocol (Nakano and Asada 1981), ascorbate peroxidase activity was assessed. Reaction mixtures in Eppendorf tubes contained 1.8 mL of 50 mM potassium phosphate buffer (pH 8.0) supplemented with 0.3 mL each of 2.5 mM EDTA, 200 mM hydrogen peroxide (H_2_O_2_), and 2.5 mM ascorbic acid. Enzymatic reactions began with the addition of 0.1 mL of enzyme extract. Absorbance changes were tracked at 290 nm using a spectrophotometer at 30-second intervals over 2 minutes. Ascorbate peroxidase activity was determined based on the rate of abs Ascorbate peroxidase activity was determined based on the rate of absorbance increase per minute.

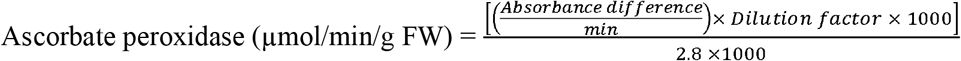

Here, Extinction Coefficient = 2.8/M/cm of APX

### Catalase Activity (CAT)

Catalase activity was evaluated following the standard methodology (Aebi 1984). In brief, a reaction mixture was prepared within an Eppendorf tube, consisting of 2.1 mL of 50 mM potassium phosphate buffer (pH 8.0), 0.3 mL of 2.5 mM EDTA solution, and 0.3 mL of 200 mM hydrogen peroxide (H_2_O_2_) solution. The enzymatic reaction commenced upon the addition of 0.1 mL of the enzyme extract. Using a spectrophotometer, changes in absorbance at 240 nm were monitored at 30-second intervals for a duration of 2 minutes. Catalase activity was subsequently determined based on the calculated increase in absorbance per minute.

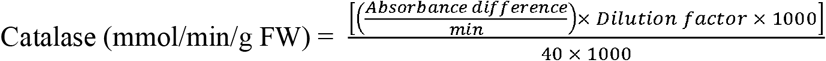

Here, Extinction Coefficient = 40/M/cm of CAT

### Statistical Analysis

A two-way analysis of variance (ANOVA) was conducted to assess the significant differences in mean performances across various parameters. Tukey’s (HSD) post-hoc test was employed for multiple comparisons among the groups. Additionally, a correlation matrix was generated using all cabbage variables to explore potential correlations. All statistical analyses were carried out using the R programming environment and visualized using MS Excel 2019.

## Results

### Antioxidant Activity, ROS, and Relative Water Content Analysis

The comparison of mean values for studied variables between red cabbage and green cabbage is presented in Figures 1 to 3. In fertilizer-treated plants, The RWC of leaf was higher in green cabbage (97.197%) than the red cabbage (97.088%) while under controlled conditions (without fertilizer) the value was lower in red cabbage (95.95%) than in green (96.58%). For the RWC of the head, a similar pattern was observed in both conditions (Figure 1).

**Figure 1.**
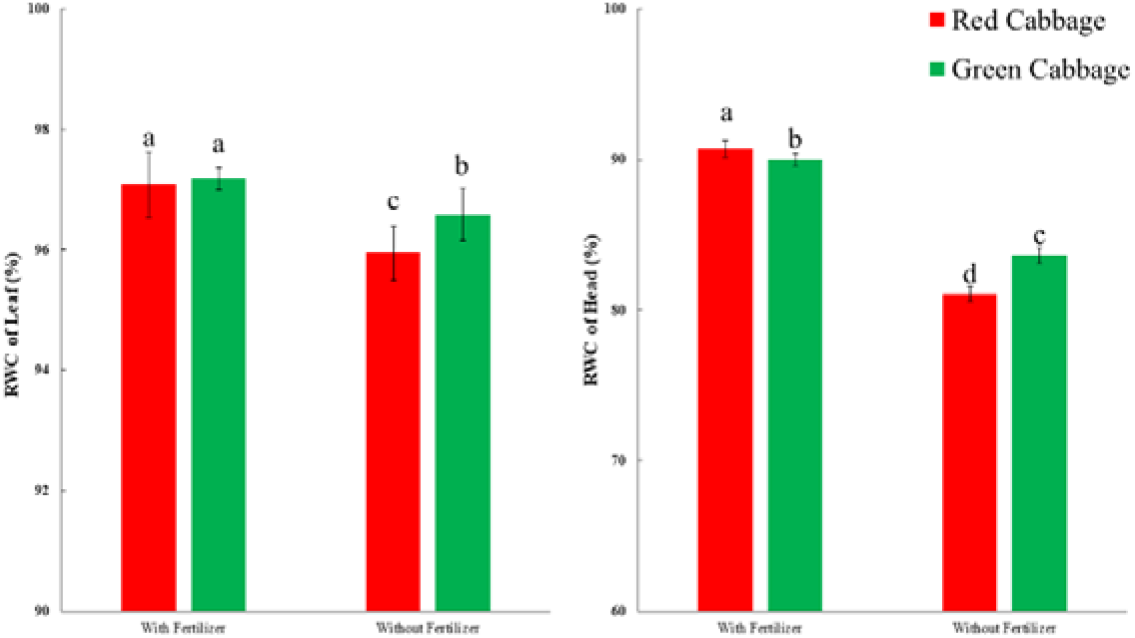
Relative Water Content (percentage) in Red and Green Cabbage. Different letters indicate statistical differences.

The MDA content was higher in red cabbage (0.242 µM/g FW) compared to its counterpart, green cabbage (0.011 µM/g FW), in fertilizer-applied plants. Similarly, in controlled conditions, the MDA content in red cabbage (0.146 µM/g FW) exceeded that of green cabbage (0.008 µM/g FW). Consistently, red cabbage exhibited greater hydrogen peroxide (H_2_O_2_) concentration compared to green cabbage under both fertilizer-treated (7.316 nM/g FW vs. 3.143 nM/g FW) and controlled condition (4.83 nM/g FW vs. 2.104 nM/g FW) conditions (Figure 2).

**Figure 2.**
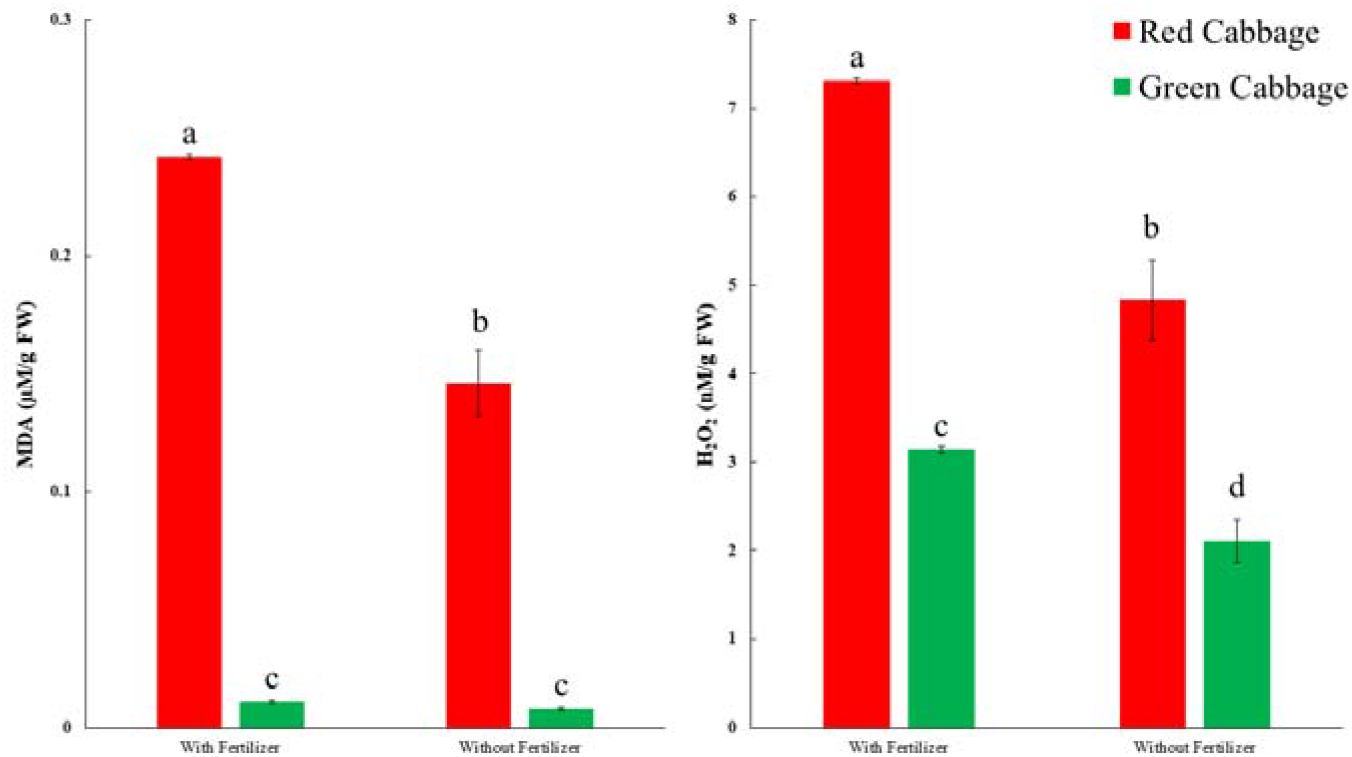
Reactive Oxygen Species (MDA and H_2_O_2_) in Red and Green Cabbage. Different letters indicate statistical differences.

**Figure 3.**
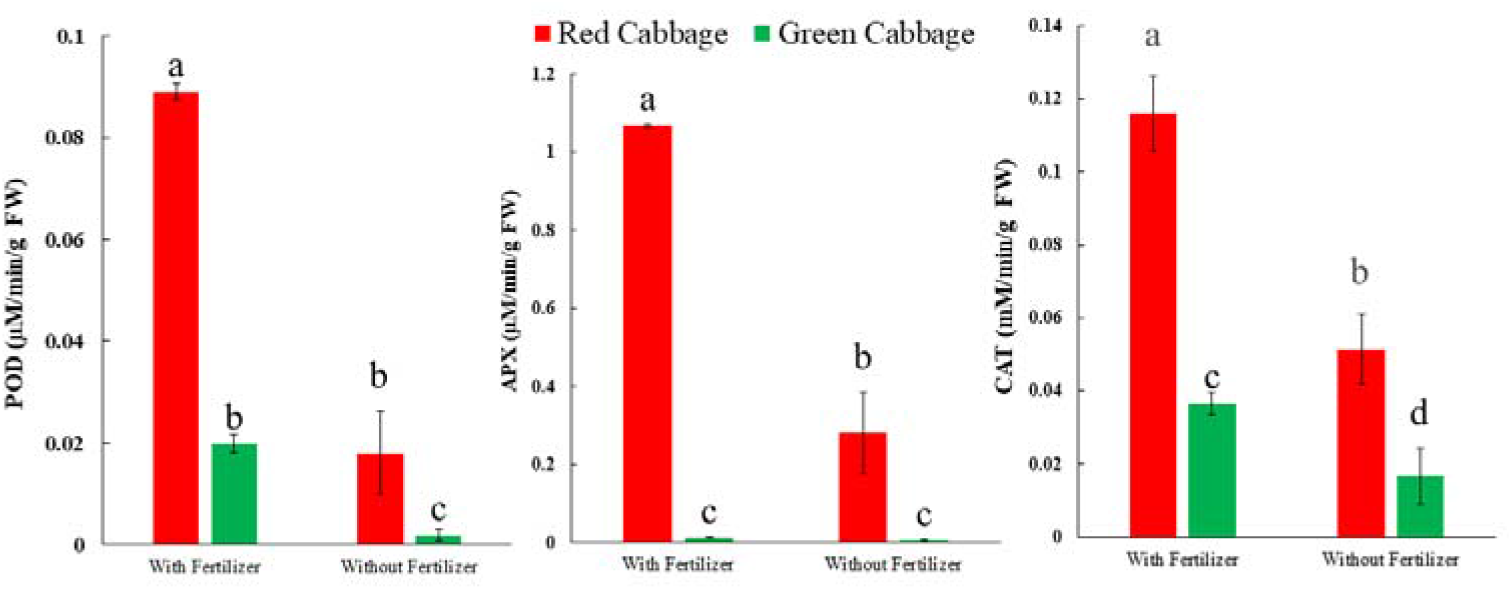
Antioxidant Activity (POD, APX, and CAT) in Red and Green Cabbage. Different letters indicate statistical differences.

Red cabbage demonstrated significantly elevated POD activity at 0.089 µM/min/g FW in contrast to green cabbage at 0.0198 µM/min/g FW when subjected to fertilizer doses. Even in controlled conditions, red cabbage maintained higher POD levels at 0.018 µM/min/g FW than green cabbage at 0.0019 µM/min/g FW. A parallel trend was observed in ascorbate peroxidase and catalase activities. Red cabbage exhibited augmented APX activity at 1.069 µM/min/g FW and 0.2812 µM/min/g FW, respectively, surpassing green cabbage values of 0.0116 µM/min/g FW and 0.0049 µM/min/g FW, in both cultivated conditions. Likewise, red cabbage displayed heightened CAT levels at 0.116 mM/min/g FW and 0.0514 mM/min/g FW compared to green cabbage at 0.0366 mM/min/g FW and 0.00167 mM/min/g FW under both experimental conditions.

### Analysis of Variance

A two-way ANOVA was carried out to understand if there were any significant variations in average performance among different parameters. For combined data, all the parameters were found to express statistically significant variations in the case of genotype, treatment, and genotype-treatment interaction at 0.1% and 1% level of significance except RWC% of leaf which showed significant variation only at 1% level.

**Table 1:** Analysis of Variance (Mean Sum Square values with the level of significance)

| Source of variations | RWC (%) |  | MDA | H<sub>2</sub>O<sub>2</sub> | POD | APX | CAT |
| --- | --- | --- | --- | --- | --- | --- | --- |
|  | Leaf | Head | 532 nm | 390 nm | 470 nm | 290 nm | 240 nm |
| Genotype | 2.072** | 13.3*** | 0.5129*** | 178.46*** | 0.027568*** | 6.673*** | 0.04937*** |
| Treatment | 11.397*** | 960.8*** | 0.0372*** | 46.61*** | 0.029221*** | 2.37*** | 0.02714*** |
| Genotype × Treatment | 1.048*** | 40.2*** | 0.0325*** | 7.84*** | 0.010399*** | 2.291*** | 0.00773*** |
| Residuals | 0.105 | 0.2 | 0.0001 | 0.07 | 0.000019 | 0.003 | 0.00007 |
| ***Significance at P<0.001, **Significance at P<0.01, *Significance at P<0.05 |  |  |  |  |  |  |  |

### Correlation Analysis

The correlation analysis for the RWC%, ROS, and antioxidant compounds was conducted separately for fertilizer treatments, and for their combined conditions (Figure 4).

**Figure 4:**
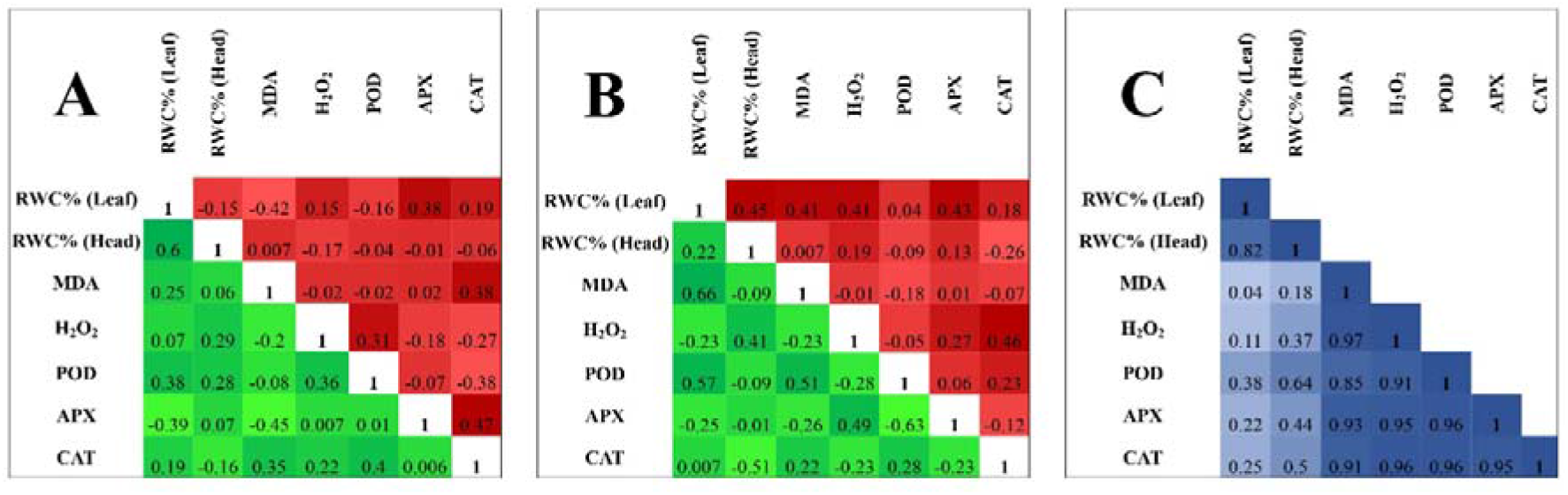
Correlation plot showing the relationship between RWC of leaf, RWC of head, MDA, H_2_O_2_, POD, APX, and CAT of green cabbage (lower matrix) and red cabbage (upper matrix) A) with fertilizer B) without fertilizer C) with and without fertilizer (combined) at α ≤ 0.05.

In fertilizer-treated plants, the analysis revealed no strong positive or negative relationships (Figure 4A). However, in green cabbage, the RWC of leaf was in moderate positive correlation with POD while in negative (moderate) correlation with APX; MDA also exhibited a moderate positive relationship with CAT but a negative one with APX. Furthermore, H_2_O_2_ and POD are positively (moderate) correlated with each other. In the case of red cabbage, the RWC of leaf was in moderate positive correlation with APX while in negative (moderate) correlation with MDA whereas MDA exhibited a moderate positive relationship with CAT. H_2_O_2_ with POD and APX with CAT are positively (moderate) correlated with each other.

There were also no strong positive or negative relationships were observed for both cabbage without fertilized conditions (Figure 4B). However, in green cabbage, the RWC of leaf was in moderate positive correlation with MDA, POD, and H_2_O_2_. Though the RWC of head was in moderately positive correlation with H_2_O_2_, it exhibited a negative one with CAT. Similarly, APX had in moderately positive correlation with H_2_O_2_, it exhibited a negative one with POD while MDA had in positive (moderate) correlation with POD. In red cabbage, the RWC of leaf was in moderate positive correlation with RWC of head, MDA, APX, and H_2_O_2_ whereas, H_2_O_2_ and CAT are positively (moderate) correlated with each other.

In overall correlation, multiple sets of significant correlations were observed at α≤0.05. Among them, a strong positive relationship between the RWC of the leaf and the RWC of the head whereas H_2_O_2,_ POD, APX, and CAT have a strong positive correlation with MDA. Similarly, H_2_O_2_ possesses a strong positive correlation with POD, APX, and CAT. Moreover, a positive association was also observed in the case of POD with APX and CAT. Furthermore, APX displayed a highly positive correlation with CAT. No negative relationships were observed in this correlation study among these variables.

## Discussions

This study provides a comprehensive comparison of the relative water content of the leaf and head and other antioxidant contents of red and green cabbage. These two types of cabbages were cultivated under both fertilizer and without fertilizer conditions. It was found that there were significant differences among the studied variables between these two varieties which is in line with several researchers (Yue et al. 2024; Zayed et al. 2023).

Green cabbage is the most common cabbage, but red cabbage has more nutritional value. Previous studies have also shown the health benefits of consuming chemoprotective substances found in red cabbage (Zayed et al. 2023; Mateljan 2001). In this study, it was also observed red cabbage possesses a higher amount of quality indices than green cabbage. Red cabbage held a higher amount of MDA, H_2_O_2_, POD, APX, and CAT content compared to green cabbage in both cultivated conditions. Red cabbage generally contains more anthocyanin pigments (Dyrby, Westergaard, and Stapelfeldt 2001), which are water-soluble and contribute to its vibrant color. These pigments are stored in the vacuoles of the cabbage cells, along with water due to this reason they contain more RWC than green. However, the relative water content of the leaf was found to be lower in red cabbage. The H_2_O_2_ content and MDA content can indicate oxidative stress levels in plant tissues, which can vary depending on factors such as environmental conditions, genotype, and developmental stage. Generally, higher levels of H_2_O_2_ and MDA suggest increased oxidative stress (Singh, Sharma, and Singh 2010). Studies have shown that red cabbage tends to have higher levels of anthocyanins compared to green cabbage. Anthocyanins are antioxidants that can help mitigate oxidative stress. In terms of, research has indicated that red cabbage has higher levels of H_2_O_2_ content compared to green cabbage due to its richer antioxidant content depending on factors such as cultivar and growing conditions (Shiyab 2015), which is in line with our findings. The rich red color of red cabbage reflects its concentration of anthocyanin polyphenols, which contribute to red cabbage containing significantly more protective phytonutrients than green cabbage. A measure of the antioxidant capacity, of red cabbage is also six to eight times higher than that of green cabbage. Mainly due to the presence of a rich amount of catalase in red one, the antioxidant activities are higher. In our findings, the catalase activity is also significantly higher in red than the green cabbage. CAT is significant in guarding against harm caused by oxidative stress by accelerating the breakdown of hydrogen peroxide into water and oxygen. By effectively neutralizing free radicals, these enzymes contribute to upholding the cellular redox balance within plant cells (Rana and Saklani 2019). The prior findings also align with our current observation (Shiyab 2015; Singh, Sharma, and Singh 2010). They also found that the CAT and APX activity is higher in red cabbages compared to green. These compounds protect biological systems against oxidative stress in the development and progression of various human diseases. Findings of our results exhibited in the comparison between green and red cabbages in terms of RWC, ROS, and antioxidants red cabbage is ahead of green cabbage, which is in line with other researchers’ findings (Zayed et al. 2023).

Knowledge of the correlation between the variables is essential as this may help in constructing suitable selection criteria for further breeding purposes (Kibar, Karaağaç, and Kar 2014). To determine the relationships between the studied variables (RWC of leaf and head, MDA, H_2_O_2,_ POD, APX, CAT) correlation coefficients were calculated, which indicates that multiple sets of associations exhibited significant positive correlations at α≤0.05.

In our study, MDA exhibited a positive correlation with H_2_O_2,_ POD, APX, and CAT. The correlation between MDA and H_2_O_2_ in cabbage is influenced by various factors. The S-methyl cysteine sulphoxide (SMCO) and thiocyanate ion (SCN−) contents in cabbage heads varied significantly, suggesting a potential impact on MDA and H_2_O_2_ levels (Bradshaw and Borzucki 2006). Higher MDA and H_2_O_2_ levels in the floral buds of a non-heading Chinese cabbage line were observed, indicating a positive correlation between these two compounds (Yuan et al. 2007). Prior studies found a positive correlation between POD and (MDA) accumulation in Brassica, with an increase in MDA under salinity stress (Hussain et al. 2024). This suggests that peroxidase activity may be linked to MDA production in response to stress. Catalase activity also showed a positive correlation with ascorbate peroxidase. Ascorbate peroxidase primarily scavenges hydrogen peroxide (H_2_O_2_) by using ascorbate as an electron donor, while catalase decomposes hydrogen peroxide into water and oxygen (Karyotou and Donaldson 2005). As the ROS levels are also found to be in a higher amount in our case the APX and CAT along with POD showed a positive association to cope with the upregulated ROS.

## Conclusions and Recommendations

This study unveils the profound influence of cultivation conditions on the relative water content, ROS, and antioxidant profile of cabbage, particularly distinguishing between red and green varieties. Notably, due to fertilizer application, both cabbages displayed enhanced leaf and head relative water content, alongside elevated MDA, H_2_O_2_, POD, APX, and CAT levels, demonstrating their resilience and antioxidant potency. These findings underscore the potential of fertilizer application to amplify the antioxidant potential of red and green cabbage, positioning it as a promising strategy for enhancing antioxidant enrichment in agricultural produce. In comparison with red and green cabbages, this study presents red cabbage as a superior source of antioxidants compared to its green counterpart, outperforming it in multiple parameters, except for leaf relative water content. This suggests that red cabbage could be preferred for those seeking to maximize antioxidant intake from cabbage consumption. Future investigations should delve into a broader context considering its breeding program and practical implications at the farmer level.

## Acknowledgements

The authors wish to express their gratitude to the Department of Agriculture at Noakhali Science and Technology University, Noakhali, Bangladesh, and the Department of Genetics and Plant Breeding at Bangladesh Agricultural University, Mymensingh, Bangladesh, for their invaluable support in conducting the field and laboratory experiments.

## Novelty Statement

This study provides novel insights into the differential impact of fertilizer application on red and green cabbage varieties, highlighting that red cabbage exhibits superior antioxidant properties and ROS levels, whereas green cabbage demonstrates higher relative water content. Moreover, the research also identifies significant variations in the nutritional properties of red and green cabbage, suggesting that red cabbage may be more beneficial for consumers seeking high antioxidant and ROS levels, thereby guiding future breeding programs to enhance these specific nutritional traits.

## Authors’ Contribution

**Md. Rayhan Sojib:** Experimental design, Field and Laboratory experiments, Data collection and analysis, Writing the manuscript (first draft)

**Nusrat Jahan Methela:** Experimental design, reviewing the manuscript, Supervising the whole experiment

**Biswajit Das and Md Nure Adil Siddique:** Writing the manuscript

**Sadia Sultana, Ataul Karim, Rafiqul Islam**: Data collection and Writing the manuscript (first draft)

**Md Jahid Hasan Jone:** Data analysis, visualization and editing of the manuscript All authors read and approved the final manuscript.

## REFERENCES

Abramowitz, C., A. Deutch, E. Shamsian, Y. Musheyev, F. Ftiha, and S. Burnstein. 2022. ‘Red Cabbage: A Novel Treatment for Periductal Lactational Mastitis’, Cureus, 14: e33191.

Aebi, H. 1984. ‘Catalase in vitro’, Methods Enzymol, 105: 121–6.

Ahmed, Shakil, Mahin Das, Md Rayhan Sojib, Shishir Kanti Talukder, Sadia Sultana, Prantika Datta, Shofiqul Islam, and Gazi Md Mohsin. 2024. ‘Evaluation of Local and Exotic Hybrid Genotypes of Yardlong Bean (Vigna unguiculata) in Saline Prone Area of Bangladesh’, Sarhad Journal of Agriculture, 40: 347–53.

Arrais, A., F. Testori, R. Calligari, V. Gianotti, M. Roncoli, A. Caramaschi, V. Todeschini, N. Massa, and E. Bona. 2022. ‘Extracts from Cabbage Leaves: Preliminary Results towards a “Universal” Highly-Performant Antibacterial and Antifungal Natural Mixture’, Biology (Basel), 11.

Ashfaq, Faiza. 2018. ‘Compositional Analysis of Pakistani Green and Red Cabbage’, Pakistan Journal of Agricultural Sciences, 55: 191–96.

Bispo, W. M. S., L. Araújo, M. B. BermúdezCCardona, I. S. Cacique, F. M. DaMatta, and F. A. Rodrigues. 2014. ‘Ceratocystis fimbriataCinduced changes in the antioxidative system of mango cultivars’, Plant Pathology, 64: 627–37.

Bradshaw, John E., and Ronald Borzucki. 2006. ‘Digestibility, SCmethyl cysteine sulphoxide content and thiocyanate ion content of cabbages for stockfeeding’, Journal of the Science of Food and Agriculture, 33: 1–5.

Braga, Daiane Cristina de Assis, Hillary Nascimento Coletro, and Maria Tereza de Freitas. 2019. ‘Nutritional composition of fad diets published on websites and blogs’, Revista de Nutrição, 32.

Chun, O. K., N. Smith, A. Sakagawa, and C. Y. Lee. 2004. ‘Antioxidant properties of raw and processed cabbages’, Int J Food Sci Nutr, 55: 191–9.

Dyrby, Marianne, Nanna Westergaard, and Henrik Stapelfeldt. 2001. ‘Light and heat sensitivity of red cabbage extract in soft drink model systems’, Food Chemistry, 72: 431–37.

FAOSTAT. 2022. ‘Crops/Regions/World list/Production Quantity (pick lists), Cabbages and other brassicas, 2020’, UN Food and Agriculture Organization, Corporate Statistical Database.

Ghareaghajlou, N., S. Hallaj-Nezhadi, and Z. Ghasempour. 2021. ‘Red cabbage anthocyanins: Stability, extraction, biological activities and applications in food systems’, Food Chem, 365: 130482.

González, Luís, and Marco González-Vilar. 2003. ‘Determination of Relative Water Content.’ in Manuel J. Reigosa Roger (ed.), Handbook of Plant Ecophysiology Techniques (Springer Netherlands: Dordrecht).

Gruszecki, Robert, Magdalena Walasek-Janusz, Gianluca Caruso, Grażyna Zawislak, Nadezhda Golubkina, Alessio Tallarita, Ewa Zalewska, and Agnieszka Sękara. 2022. ‘Cabbage in Polish folk and veterinary medicine’, South African Journal of Botany, 149: 435–45.

Hussain, Saber, Shakil Ahmed, Waheed Akram, Aqeel Ahmad, Nasim Ahmad Yasin, Mei Fu, Guihua Li, and Rehana Sardar. 2024. ‘The potential of selenium to induce salt stress tolerance in Brassica rapa: Evaluation of biochemical, physiological and molecular phenomenon’, Plant Stress, 11: 100331.

Islam, Nadia, Md Hashan, Rayhan Ahammed, Biswajit Das, Mst Naher, and Shohrab Hoshain. 2023. ‘Effect of Various Doses of Cowdung and Nitrogen on The Yield Performance of Mustard in Coastal Area of Bangladesh (Brassica Sp.)’, Research in Agriculture, Livestock and Fisheries, 10: 109–22.

Karyotou, K., and R. P. Donaldson. 2005. ‘Ascorbate peroxidase, a scavenger of hydrogen peroxide in glyoxysomal membranes’, Arch Biochem Biophys, 434: 248–57.

Kasarello, K., I. Kohling, A. Kosowska, K. Pucia, A. Lukasik, A. Cudnoch-Jedrzejewska, L. Paczek, U. Zielenkiewicz, and P. Zielenkiewicz. 2022. ‘The Anti-Inflammatory Effect of Cabbage Leaves Explained by the Influence of bol-miRNA172a on FAN Expression’, Front Pharmacol, 13: 846830.

Kibar, Beyhan, Onur Karaağaç, and Hayati Kar. 2014. ‘Correlation and path coefficient analysis of yield and yield components in cabbage (Brassica oleracea var. capitata L.)’, Acta Scientiarum Polonorum, Hortorum Cultus, 13: 87–97.

Liang, Ying, Yi Li, Liujuan Zhang, and Xianjin Liu. 2019. ‘Phytochemicals and antioxidant activity in four varieties of head cabbages commonly consumed in China’, Food Production, Processing and Nutrition, 1: 3.

Manchali, Shivapriya, Kotamballi N. Chidambara Murthy, and Bhimanagouda S. Patil. 2012. ‘Crucial facts about health benefits of popular cruciferous vegetables’, Journal of Functional Foods, 4: 94–106.

Mateljan, George. 2001. The world’s healthiest foods (The George Mateljan Foundation).

Melim, C., M. R. Lauro, I. M. Pires, P. J. Oliveira, and C. Cabral. 2022. ‘The Role of Glucosinolates from Cruciferous Vegetables (Brassicaceae) in Gastrointestinal Cancers: From Prevention to Therapeutics’, Pharmaceutics, 14.

Misu, Israt Jahan, Ali Md Sabuj, Rony Ashraful Islam, Siddike Md Nomun, Payel Naushin Alim, Islam Md Fakhrul, Tanvir Shohel, Shikder Md Mafin, and Mohsin Gazi Md. 2023. ‘Effect of plant growth regulators on growth, yield and yield contributing characters of brinjal (Solanum melongena L.) in coastal zone of Bangladesh’, Research in Agriculture Livestock and Fisheries, 10: 91–97.

Moreb, Nora, Amy Murphy, Swarna Jaiswal, and Amit K. Jaiswal. 2020. ‘Cabbage.’ in, Nutritional Composition and Antioxidant Properties of Fruits and Vegetables.

Nakano, Yoshiyuki, and Kozi Asada. 1981. ‘Hydrogen Peroxide is Scavenged by Ascorbate-specific Peroxidase in Spinach Chloroplasts’, Plant and Cell Physiology, 22: 867–80.

Osman, Muhammad, Kawsar Hossen, Rabiul Chowdhury, Chowdhury Tabassum, Khairul Islam, Rayhan Ahmed, and Tahmina Ferdous. 2020. ‘Assessment of Different Weed Control Methods on Growth and Yield Performance of T. Aus Rice’, Agricultural Research & Technology: Open Access Journal, 24.

Pérez, Amando, and Maria Cartea. 2008. ‘Cabbage and Kale’, 1.

Rana, Pawan Singh, and Pooja Saklani. 2019. ‘ANALYZING THE EFFECT OF ALTITUDINAL VARIATION IN ENZYMATIC ANTIOXIDANTS OF Coleus forskohlii FROM UTTARAKHAND, INDIA’, Plant Cell Biotechnology and Molecular Biology, 20: 442–50.

Šamec, Dunja, Jasenka Piljac-Žegarac, Mara Bogović, Ksenija Habjanic, and Jiří Grúz. 2011. ‘Antioxidant potency of white (Brassica oleracea L. var. capitata) and Chinese (Brassica rapa L. var. pekinensis (Lour.)) cabbage: The influence of development stage, cultivar choice and seed selection’, Scientia Horticulturae, 128: 78–83.

Shiyab, Safwan. 2015. ‘Impact of cadmium accumulation on physiological characteristics of two cabbage varieties during phytoremediation’, Journal of Food, Agriculture and Environment, 13: 262–68.

Singh, B. K., S. R. Sharma, and B. Singh. 2010. ‘Antioxidant enzymes in cabbage: Variability and inheritance of superoxide dismutase, peroxidase and catalase’, Scientia Horticulturae, 124: 9–13.

Singh, Jagdish, A. K. Upadhyay, A. Bahadur, B. Singh, K. P. Singh, and Mathura Rai. 2006. ‘Antioxidant phytochemicals in cabbage (Brassica oleracea L. var. capitata)’, Scientia Horticulturae, 108: 233–37.

Tsania, Nidya Ulfana, Sandhika Willy, and Sawitri. 2023. ‘The Effect of Cabbage (Brassica oleracea var. capitata L.) Extract on Macrophage and Blood Vessel Counts in Clean Wound Tissue of Male Rats (Rattus norvegicus)’, Folia Medica Indonesiana, 59: 136–42.

Turner, E. R., Y. Luo, and R. L. Buchanan. 2020. ‘Microgreen nutrition, food safety, and shelf life: A review’, J Food Sci, 85: 870–82.

Vega-Galvez, A., L. S. Gomez-Perez, F. Zepeda, R. L. Vidal, F. Grunenwald, N. Mejias, A. Pasten, M. Araya, and K. S. Ah-Hen. 2023. ‘Assessment of Bio-Compounds Content, Antioxidant Activity, and Neuroprotective Effect of Red Cabbage (Brassica oleracea var. Capitata rubra) Processed by Convective Drying at Different Temperatures’, Antioxidants (Basel), 12.

Yuan, Jianyu, Xilin Hou, Changwei Zhang, and Fan Ye. 2007. ‘Active oxygen metabolism in the floral buds and leaves of the new cytoplasm male sterile (CMS) line and its maintainer line of non-heading Chinese cabbage’, Frontiers of Agriculture in China, 1: 47–51.

Yue, Z., G. Zhang, J. Wang, J. Wang, S. Luo, B. Zhang, Z. Li, and Z. Liu. 2024. ‘Comparative study of the quality indices, antioxidant substances, and mineral elements in different forms of cabbage’, BMC Plant Biol, 24: 187.

Zayed, A., M. Sheashea, I. A. A. Kassem, and M. A. Farag. 2023. ‘Red and white cabbages: An updated comparative review of bioactives, extraction methods, processing practices, and health benefits’, Crit Rev Food Sci Nutr, 63: 7025–42.

